# Compositionally unique mitochondria in filopodia support cellular migration

**DOI:** 10.1101/2024.06.21.600105

**Authors:** Madeleine Marlar-Pavey, Daniel Tapias-Gomez, Marcel Mettlen, Jonathan R. Friedman

## Abstract

Local metabolic demand within cells varies widely and the extent to which individual mitochondria can be specialized to meet these functional needs is unclear. We examined the subcellular distribution of MICOS, a spatial and functional organizer of mitochondria, and discovered that it dynamically enriches at the tip of a minor population of mitochondria in the cell periphery that we term “METEORs”. METEORs have a unique composition; MICOS enrichment sites are depleted of mtDNA and matrix proteins and contain high levels of the Ca^2+^ uniporter MCU, suggesting a functional specialization. METEORs are also enriched for the myosin MYO19, which promotes their trafficking to a small subset of filopodia. We identify a positive correlation between the length of filopodia and the presence of METEORs and show that elimination of mitochondria from filopodia impairs cellular motility. Our data reveal a novel type of mitochondrial heterogeneity and suggest compositionally specialized mitochondria support cell migration.

## Introduction

Mitochondria are double membrane-bound organelles responsible for energy production and numerous other critical cellular functions, including metabolite transport, lipid synthesis, and the maintenance of ion homeostasis (*1, 2*). Mitochondria form a semi-continuous network whose shape and subcellular distribution is controlled via cytoskeleton-based motility, division, and homotypic fusion (*3, 4*). Due to these fission and fusion dynamics, mitochondria can exist as a single organelle, or up to hundreds or thousands of individual organelles within cells. Individual mitochondria can also be highly heterogeneous, having variable protein density and composition, shapes, inter-organelle contacts, and functions (*5–11*). However, we have a minimal understanding of the extent to which individual mitochondria play specific roles in response to local metabolic needs within cells, and how this physical and functional heterogeneity is established.

One of the key contributors to mitochondrial function is the elaborate organization of the inner mitochondrial membrane (IMM) into three morphologically distinct domains (*12*). The IMM is closely apposed to the outer mitochondrial membrane (OMM) at the boundary, a domain which is responsible for protein import and trafficking between the inside and outside of mitochondria. The IMM invaginates from the boundary to form cristae membranes, which house respiratory complexes responsible for ATP synthesis. The boundary and cristae domains are connected by cristae junctions, narrow tubular necks that are thought to organize the domains and prevent the free diffusion of proteins between each IMM sub-compartment (*13*). Mitochondrial cristae density correlates with the respiratory function of the organelle, and the IMM can dynamically remodel to increase cristae number in response to elevated respiratory demand (*12, 14*). The IMM can also re-organize to serve other roles, including the release of cytochrome *c* to mediate apoptosis (*15*). The number and organization of cristae membranes is established in large part by the Mitochondrial Contact Site and Cristae Organizing System (MICOS) complex, which forms a ∼megadalton-sized assembly that enriches at cristae junctions (*16*). MICOS has membrane shaping roles and is thought to provide stability to the curvature of the cristae junction (*17–19*). Disruption of MICOS leads to a near complete loss of cristae junctions, severely affects mitochondrial cristae morphology, and is linked to numerous pathologies, including Parkinson’s Disease, diabetes, and aging (*20–22*). In addition to its role in cristae organization, MICOS is a “master regulator” of mitochondrial function, forming an extensive protein interaction network with protein complexes and machineries on the IMM and OMM (*23–29*). Among these interactions, MICOS forms a supercomplex, the mitochondrial intermembrane space bridging complex (MIB), with OMM protein biogenesis machinery (*30, 31*). Thus, in addition to its role as a key determinant of cristae architecture and respiratory function, MICOS is central to overall mitochondrial organization.

Here, we describe a distinct subpopulation of mitochondria in the cell periphery that display a high degree of MICOS enrichment at their distal tips. Based on the <u>M</u>ICOS <u>e</u>nrichment at the tips of mitochondria, and their distinctive appearance, we refer to these mitochondria hereafter as “METEORs”. Using a candidate-based immunolabeling approach, we characterize METEORs as having a highly unique composition. While they maintain membrane potential and contain respiratory complexes, MICOS enrichment sites are generally de-enriched for matrix proteins and absent of mtDNA. Further, they are enriched for the Ca^2+^ uniporter MCU. We also determine that METEORs traffic dynamically into a subset of filopodia, thin actin-mediated plasma membrane (PM) extensions, in a manner dependent on the OMM-associated myosin MYO19. Finally, using two distinct approaches, we prevent METEOR formation and trafficking into filopodia, both of which lead to a decrease in filopodia length and reduced cellular motility. Together, our data reveal that MYO19 and the MICOS complex help establish a compositionally unique subpopulation of mitochondria that reside within filopodia and positively support cellular migration.

## Results

### MICOS complexes are highly enriched at the tip of a subset of mitochondria in the cell periphery

Previously, we performed indirect immunofluorescence of the MICOS subunit MIC60 in U2OS cells, an osteosarcoma cell line (*28, 32*). In most mitochondria, MICOS is distributed in an uneven semi-punctate pattern, corresponding to the enrichment of the protein at cristae junctions (*33, 34*). However, we noticed that in some cells, a small number of mitochondria in the cellular periphery appear highly enriched for MIC60. To examine this phenomenon in more detail, we immunolabeled and imaged MIC60 and TIMM23, a member of the translocase of the inner membrane (TIM) complex and an abundant IMM protein (*35, 36*). Strikingly, while MIC60 and TIMM23 had a similar distribution in most mitochondria, MIC60 occasionally appeared highly concentrated relative to TIMM23 at the distal tip of a mitochondrion (Fig. 1A, see linescan), or enriched in a small mitochondrial fragment, in the cell periphery. These mitochondria with MIC60-enriched tips (METEORs) are present in about 40% of cells, usually as three or fewer mitochondria per cell (Fig. 1B). To determine if MIC60 is enriched specifically compared to TIMM23, or other proteins as well, we immunolabelled cells with the translocase of the outer membrane (TOM) subunit TOMM20 (*36*). Similar to TIMM23, we observed examples of mitochondria in the cell periphery with local enrichment of MIC60 with no corresponding increase in TOMM20 signal (Fig. 1C). Notably, we observed TOMM20-labeled cells with both METEORs and mitochondrial derived vesicles (MDVs), indicating that the MIC60-enriched METEORs are distinct from MDVs (*37, 38*) (Fig. S1A).

**Figure 1.**
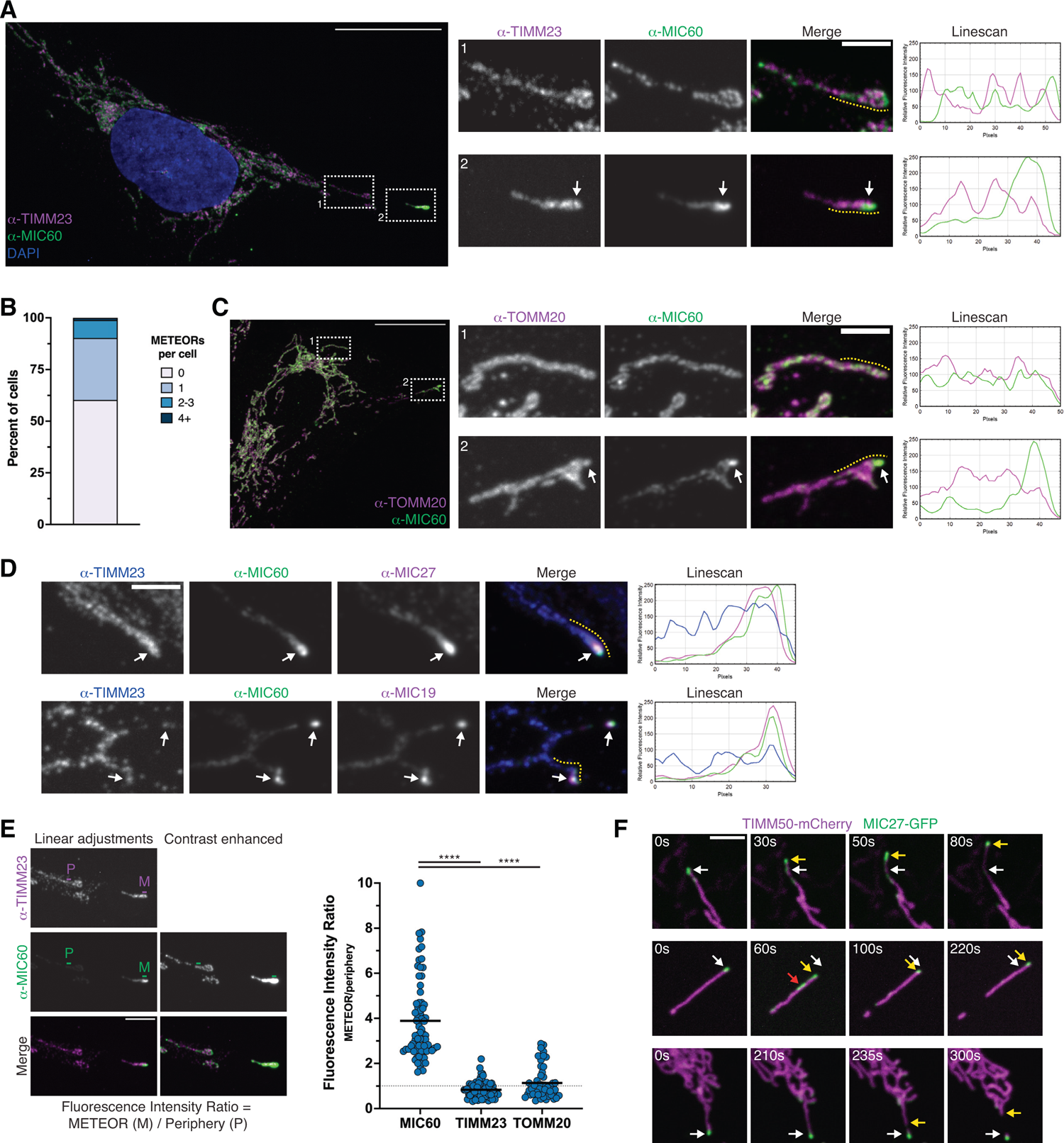
MICOS complexes are highly enriched at the tip of a subset of mitochondria in the cell periphery. **(A)** Maximum intensity projection confocal images are shown of a fixed U2OS cell immunolabeled for TIMM23 and MIC60, and stained with DAPI. Dotted boxes correspond to the numbered insets shown on the right. Box 1 shows a representative mitochondrion in the cell periphery and Box 2 shows a METEOR. The yellow dotted lines on the insets correspond to the fluorescence intensity linescans shown at the right. White arrows mark a site of MIC60 enrichment. **(B)** A graph depicting the frequency of METEORs from cells labeled as in (A) with a 2-fold or higher relative enrichment of MIC60. See (E) for an explanation of quantitative analysis. Data shown represent a total of 174 cells from two independent experiments. **(C)** Images are shown as in (A) for a cell immunolabeled for TOMM20 and MIC60. **(D)** As in (A) for a cell immunolabeled for TIMM23, MIC60, and either MIC27 (top panel) or MIC19 (bottom panel). **(E)** Left: A schematic is shown detailing the quantitative analysis used to determine the ratio of the fluorescence intensity at the tip of METEORs (M) relative to mitochondria in the cell periphery (P). See Methods for additional details. The example image is redisplayed from (A). Right: A graph of calculated fluorescence intensity ratios for MIC60 (n=71), TIMM23 (n=76), and TOMM20 (n=49). Data were acquired from two independent experiments. Solid lines indicate means, and a dotted line is shown at y=1 for reference. **(F)** Single plane live cell confocal images are shown of METEORs at the indicated timepoints (s = seconds) from cells transiently transfected with TIMM50-mCherry and MIC27-GFP. The top panel depicts stable MIC27 enrichment within a dynamic mitochondrion. The white arrows mark the original MICOS enrichment position and yellow arrows track its movements. The middle panel depicts a METEOR where MIC27 dynamically redistributes away from (red arrow) and towards the mitochondrial tip (yellow arrow). The bottom panel depicts a METEOR which undergoes a fission event (yellow arrow) at the site of MICOS enrichment (white arrow). See also Fig. S1D for corresponding whole cell and unmerged images and Videos 1-3. Asterisks (****p<0.0001) represent an ordinary one-way ANOVA with Bonferroni’s multiple comparison’s test. Scale bars: (A, C) 20 µm (3 µm in inset); (D, F) 3 µm; (E) 5 µm. See also Figure S1.

To our knowledge, this distinct MIC60 distribution pattern has not been reported previously. We considered that it could be an artifact of the MIC60 antibody or not otherwise generalizable to the holo-MICOS complex. To address this, we immunolabeled cells with TIMM23, MIC60, and the MICOS subunits MIC19 or MIC27, which are members of two distinct subcomplexes (*16*). In each case, we observed co-enrichment of MIC27 and MIC19 with MIC60 relative to TIMM23 at the distal tips of mitochondria (Fig. 1D). MICOS is part of the MIB supercomplex with OMM proteins, including the Sorting and Assembly Machinery (SAM) complex and DNAJC11 (*30, 31*). Both the SAM subunit SAMM50 and DNAJC11 were also co-enriched with MIC60 at these mitochondria (Fig. S1B-S1C). Thus, components of both the MICOS and MIB complexes enrich at the tips of METEORs.

We next quantified MIC60 enrichment at the tip of METEORs. Using staining of another MICOS subunit as a proxy to blindly identify METEORs based on qualitative appearance, we determined the relative fluorescence intensity of MIC60, TIMM23, and TOMM20 at METEOR tips compared to nearby mitochondria in the cell periphery (see Methods for details). While TIMM23 and TOMM20 labeled MICOS-enriched mitochondria to a similar extent as other cell peripheral mitochondria (mean fluorescence intensity ratio of 0.8 and 1.1, respectively), MIC60 was enriched 3.9-fold on average at METEOR tips (Fig. 1E).

To exclude artifacts caused by chemical fixation and staining, we examined MICOS distribution in live cells. We transiently transfected cells with MIC27-GFP or MIC10-GFP along with the IMM marker and TIM subunit TIMM50-mCherry (*39*). When expressed at low levels, MIC27-GFP and MIC10-GFP localized to mitochondria in a patchy distribution, resembling immunolabeling of endogenous MICOS subunits (Fig. S1D-S1E). We also frequently saw examples of mitochondria at the periphery of cells with enrichments of MIC27-GFP or MIC10-GFP relative to the IMM marker TIMM50-mCherry (Fig. 1F, Fig. S1D-S1E). Next, to characterize the dynamic behavior of the MICOS subunits at these sites, we performed timelapse imaging. The localization of MICOS at the tips of METEORs often appeared stable, maintaining its relative enrichment over the course of five minutes (Fig. 1F, top panel and Video 1; Fig. S1E). However, we could also observe dynamic behavior of METEORs, including some redistribution of MIC27-GFP away from and returning to the mitochondrial tip (Fig. 1F, middle panel and Video 2). METEORs were also frequently motile (Fig. 1F, top two panels) and occasionally underwent fission near the site of MICOS enrichment to generate MICOS-enriched mitochondrial fragments (Fig. 1F, bottom panel and Video 3).

### METEORs localize to a subset of filopodia

We noticed that METEORs are found consistently at the extreme periphery of the mitochondrial network and therefore wanted to examine their localization relative to the PM. We performed post-fixation labeling of the PM with wheat germ agglutinin (WGA) conjugated to a fluorescent dye, followed by immunolabeling for MIC60 and TIMM23. Interestingly, METEORs nearly always appeared to contact the PM at the resolution of light microscopy, or localize within extensions from the PM (Fig. 2A-2B, see arrows). To visualize the dynamics of these mitochondria in live cells, we co-transfected cells with MIC27-GFP, TIMM50-mCherry, and a marker of the PM (PM-TagBFP; (*40*)), and imaged by confocal microscopy for a period of five minutes. METEORs often dynamically moved within PM extensions, including toward or with the tip of growing PM extensions (Fig. 2C, top panel and Video 4). However, mitochondria did not universally move outward, and we could see examples of mitochondria retracting from the tip of PM extensions (Fig. 2C, bottom panel and Video 5). Based on their appearance, we considered the PM extensions that contain mitochondria could be filopodia, thin actin-mediated cell protrusions that play a role in cell adhesion, environment sensing, and migration (*41, 42*). We therefore transiently transfected cells with MIC27-mCherry, PM-TagBFP, and GFP-MYO10, a marker of filopodia (*43*). The tips of PM extensions containing METEORs were routinely labeled with GFP-MYO10, indicating that these structures are indeed filopodia (Fig. 2D).

**Figure 2.**
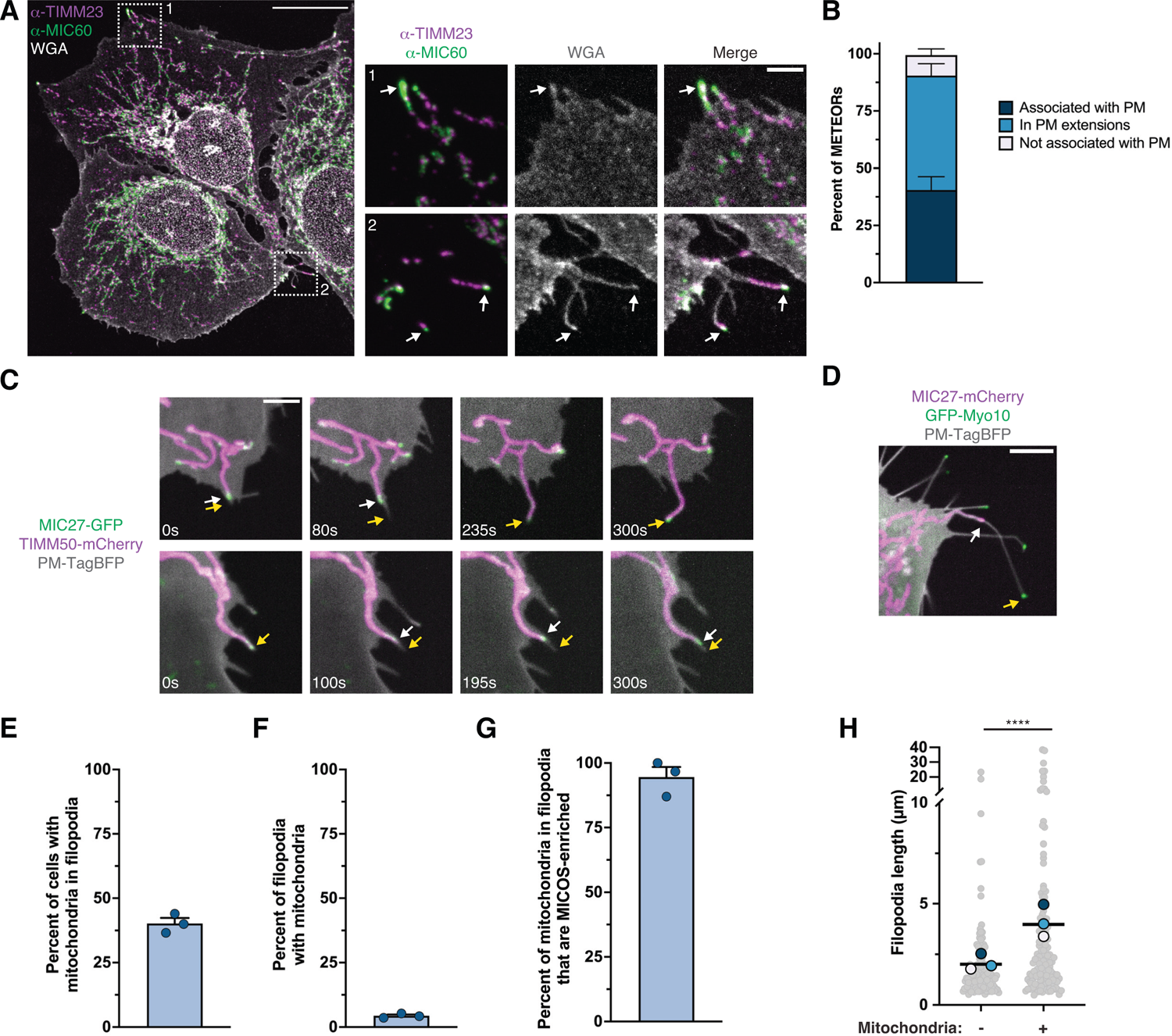
METEORs localize to a subpopulation of filopodia. **(A)** Maximum intensity projection confocal images are shown of a fixed U2OS cell immunolabeled for TIMM23 and MIC60 and stained with fluorescence-conjugated WGA. Dotted boxes correspond to the numbered insets shown on the right. Arrows correspond to METEORs in PM extensions. **(B)** A graph depicting the frequency of METEOR localization relative to the PM from cells labeled as in (A). Data shown represent >50 METEORs in each of three independent experiments and bars indicate S.E.M. METEORs were counted as “in proximity” to the PM if they appeared to be touching the PM at the resolution of light microscopy. **(C)** Single plane live cell confocal images are shown of METEORs at the indicated timepoints from cells transiently transfected with MIC27-GFP, TIMM50-mCherry, and PM-TagBFP. METEORs (white arrows) dynamically move towards the tip of a PM extension (yellow arrow; top panel) and retract from the tip of a PM extension (bottom panel). See Videos 4-5. **(D)** A single-plane confocal image is shown of a cell transiently transfected with MIC27-mCherry, GFP-MYO10, and PM-TagBFP. The white arrow indicates a METEOR in a filopodium labeled with GFP-MYO10 (yellow arrow). **(E)** A graph depicting the frequency of cells containing mitochondria in filopodia from cells labeled as in (A). Data shown represent >50 cells in each of three independent experiments and bars indicate S.E.M. **(F)** A graph depicting the frequency of filopodia containing mitochondria from cells labeled as in (A). Data shown represent >400 filopodia in each of three independent experiments and bars indicate S.E.M. **(G)** A graph depicting the frequency of mitochondria in filopodia that are METEORs from cells labeled as in (A). Data shown represent >18 filopodia-localized mitochondria in each of three independent experiments and bars indicate S.E.M. **(H)** A graph depicting the individual lengths of filopodia that contain mitochondria, or adjacent filopodia without mitochondria, from cells labeled as in (A). Data shown represent n=134 total filopodia per category acquired from three independent experiments. Solid lines indicate the mean filopodia length. Superimposed colored circles represent the means for individual experimental replicates. Asterisks (****p<0.0001) represent a paired two-tailed *t* test. Scale bars: (A) 20 µm (3 µm in inset); (C) 3 µm; (D) 5 µm.

To further characterize filopodia and their relationship to METEORs, we analyzed wild-type cells that were fixed, stained with fluorescence-conjugated WGA, and immunolabeled for MIC60 and TIMM23. Filopodia were present in nearly all cells (95.4%; n>40 cells in each of three independent experiments), and we could detect TIMM23-labeled mitochondria in at least one filopodia in ∼40% of cells (Fig. 2E). However, the presence of mitochondria within filopodia was relatively infrequent, and only 4% of all filopodia contained a mitochondrion (Fig. 2F). Next, to determine whether the presence of mitochondria within filopodia was an indicator of MICOS-enrichment, we quantitatively assessed their MIC60 staining relative to other mitochondria in the cell periphery. Strikingly, ∼95% of mitochondria within filopodia exhibited at least 2-fold enrichment of MIC60, indicating that mitochondria in filopodia are almost exclusively METEORs (Fig. 2G).

Given that only a small subset of filopodia contained mitochondria, we next asked if there were defining differences of filopodia that contain mitochondria. We compared the length of filopodia that contain mitochondria to adjacent filopodia without a mitochondria present, finding that mitochondria-containing filopodia were on average about twice as long (4.0 µm vs. 2.0 µm, respectively; Fig. 2H). Together, these data indicate that in steady-state growth conditions, a small subset of filopodia contain METEORs, the presence of which positively correlates with filopodia length.

### MYO19 and MIC60 are required for mitochondrial trafficking into filopodia

Based on our observation that METEORs were present in filopodia, we studied the mechanism by which these mitochondria were trafficked to the cellular periphery. Previous reports suggest that a mitochondrial-associated myosin, MYO19, promotes mitochondrial targeting to filopodia and filopodia formation during acute stress, including glucose starvation and H_2_O_2_ treatment (*44, 45*). Recently, MYO19 was also shown to associate with the MIB complex, possibly via a direct interaction with the SAM complex subunit MTX3 (*46, 47*). Therefore, we reasoned that MYO19 could be responsible for the targeting of mitochondria to filopodia that we observe during basal growth conditions. To explore a potential connection between MYO19 and the trafficking of METEORs to filopodia, we first examined the localization of MYO19 relative to MICOS. We fixed and immunolabeled cells for MYO19, MIC60, and TIMM23 and imaged by confocal microscopy. Consistent with previous reports (*47, 48*), immunolabeled MYO19 decorated the exterior of typical mitochondria (Fig. 3A, top panel). However, in METEORs, MYO19 concentrated at punctate structures coincident with MIC60 enrichment sites (6.7-fold mean enrichment of MYO19; Fig. 3A, bottom panel and Fig. 3B).

**Figure 3.**
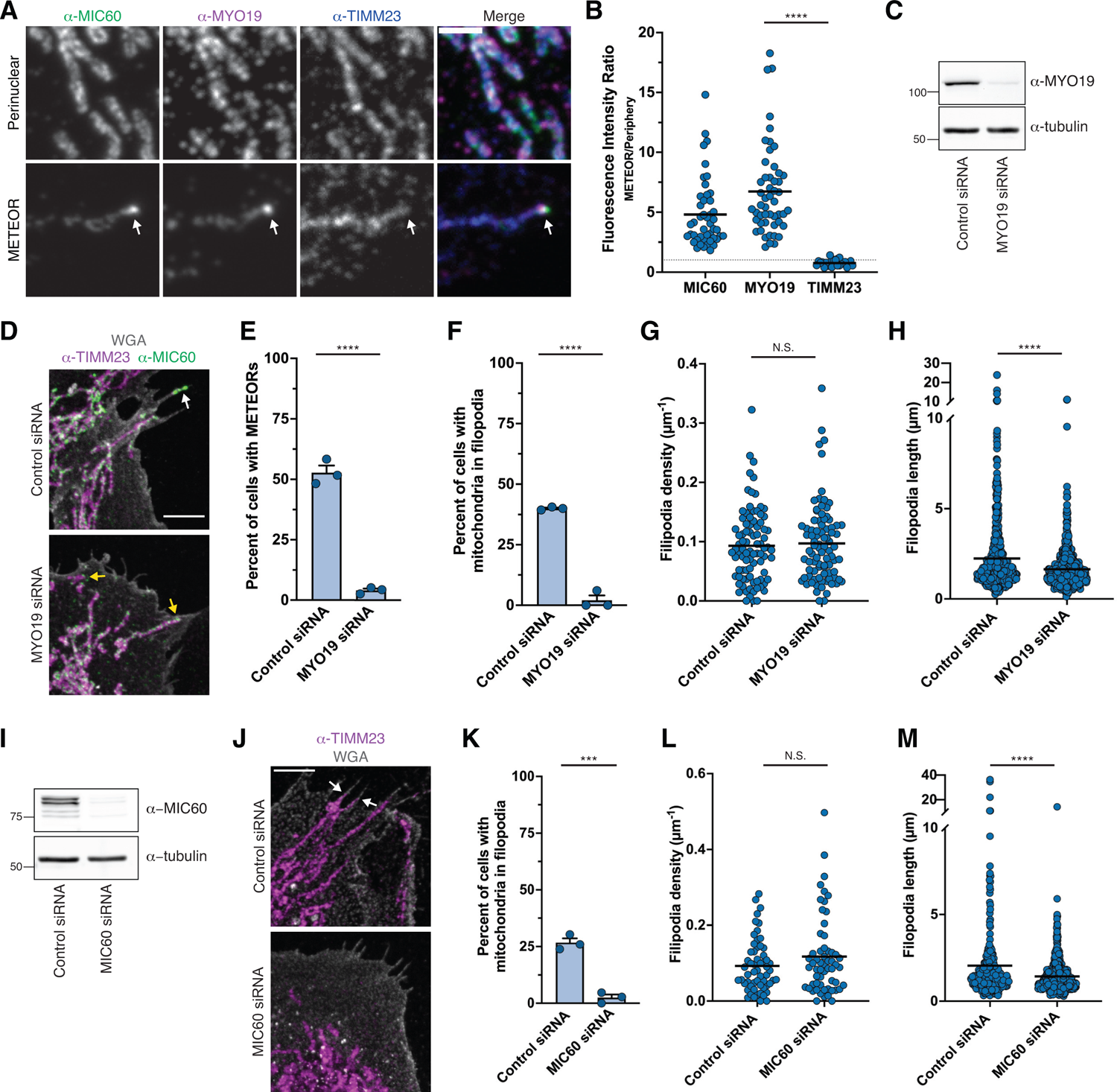
MYO19 and MIC60 are required for mitochondrial trafficking into filopodia. **(A)** Maximum intensity projection confocal images are shown of a fixed U2OS cell immunolabeled for MIC60, MYO19, and TIMM23. Images from the top row are from representative perinuclear mitochondria and images at bottom are from a METEOR in the cell periphery. White arrows mark a site of MYO19 co-enrichment with MIC60 at the tip of a METEOR. **(B)** A graph is shown of the fluorescence intensity ratios of MIC60 (n=46), MYO19 (n=51), and TIMM23 (n=26) at METEOR tips relative to nearby mitochondria in the cell periphery. Solid lines indicate the mean, and a dotted line is shown at y=1 for reference. **(C)** A Western blot with the indicated antibodies of lysate from cells transfected with a scrambled control siRNA or MYO19-targeting siRNA #1. **(D)** Representative maximum projection images of cells treated with siRNA as in (C) and fixed and immunolabeled for TIMM23, MIC60, and stained with fluorescence-conjugated WGA. The white arrow marks a METEOR localized in a filopodium. Yellow arrows in the MYO19-depleted cell highlight mitochondria localized to the cell periphery, but not localized to filopodia or enriched for MIC60. **(E-F)** Graphs depicting the frequency of cells with METEORs (E) and mitochondria in filopodia (F) in control or MYO19 siRNA treated cells as in (C). Data shown represent >30 cells per condition in each of three independent experiments, and bars indicate S.E.M. **(G)** A graph depicting the density of filopodia in individual control and MYO19 siRNA treated cells as in (C). Data represent a total of n=87 cells per condition acquired from three independent experiments. **(H)** A graph is shown depicting the length of filopodia in control or MYO19 siRNA treated cells as in (C). Data shown represent approximately 600 filopodia per condition acquired from three independent experiments. **(I)** As in (C) for cells treated with control or MIC60-targeting siRNA. **(J)** Maximum projection images of cells treated as in (I) and fixed and immunolabeled for TIMM23 and stained with fluorescence-conjugated WGA. White arrows correspond to mitochondria in filopodia. **(K)** As in (F) for cells treated as in (I). Data shown represent >30 cells per condition in each of three independent experiments and bars indicate S.E.M. **(L)** As in (G) for cells treated as in (I). Data shown represent a total of n=58 cells per condition acquired from three independent experiments. **(M)** As in (H) for cells treated as in (I). Data shown represent >300 filopodia per condition acquired from two independent experiments. Asterisks (****p<0.0001, ***p<0.001) represent an ordinary one-way ANOVA with Bonferroni’s multiple comparisons test (B) or unpaired two-tailed *t* tests (E, F, G, H, K, L, M). N.S. indicates not statistically significant. Scale bars: (A) 2 µm; (D,J) 5 µm. See also Figure S2.

We next asked whether MYO19 was required for the formation of METEORs or the basal targeting of mitochondria to filopodia. We performed transient knockdowns of MYO19 with two independent siRNAs, leading in each case to a robust reduction in MYO19 protein levels as assessed by Western blotting (Fig. 3C and Fig. S2A). We then fixed the cells, stained the PM, and immunolabeled for MIC60 and TIMM23. While MIC60 protein levels and its distribution in a semi-punctate pattern on most mitochondria was not affected by MYO19 depletion, we observed a near complete loss of MIC60-enriched METEORs (Fig. 3D-3E and Fig. S2B-S2C). In addition, we found that mitochondria were nearly entirely absent from filopodia in cells depleted of MYO19 (Fig. 3F and Fig. S2D). Together, these data indicate that MYO19 is required not only for METEOR formation, but also for trafficking of mitochondria into filopodia.

Because mitochondria were no longer present in filopodia, we next examined the consequence of MYO19 depletion on the number of filopodia and their length. Importantly, MYO19 depletion did not significantly alter the density of filopodia (Fig. 3G). However, consistent with previous observations in stressed cells (*45*), filopodia were significantly shorter in the absence of MYO19 (Fig. 3H). Notably, a small population of longer filopodia appeared selectively diminished in MYO19-depleted cells. In combination with our observation of the presence of mitochondria in longer filopodia (Fig. 2H), these data further suggest MYO19-dependent mitochondrial targeting is required to promote formation of a subset of longer filopodia during basal growth conditions.

MYO19 has been implicated in mitochondrial cristae architecture due to its interaction with MICOS/MIB, and has also has been suggested to influence mitochondrial fission dynamics and inheritance during cell division (*46, 47, 49, 50*). We therefore assessed mitochondrial morphology in cells transiently depleted of MYO19. We found that mitochondria were primarily absent from filopodia but remained in proximity to the PM (Fig. S2E-S2F). In addition, overall mitochondrial morphology as assessed by fluorescence microscopy was not grossly impacted by MYO19 depletion (Fig. S2G). We then examined mitochondrial ultrastructure by electron microscopy. In contrast to reports in MYO19 knockout cells (*46*), mitochondrial cristae appearance and density were unaffected in cells transiently depleted of MYO19 (Fig. S2H-S2I). Together, these data suggest that acute MYO19 depletion selectively decreases METEOR formation and mitochondrial targeting to filopodia without causing gross defects in the MICOS complex or mitochondrial morphology.

We next determined whether enrichment of the MICOS complex is important for the targeting of mitochondria to filopodia. We depleted the core MICOS subunit MIC60 by siRNA, which causes destabilization of the holo-MICOS complex (*34, 51*), and performed post-fixation labeling of the PM and immunolabeling of TIMM23. Consistent with our previous observations, depletion of MIC60 in U2OS cells caused mitochondrial morphology defects, including mitochondrial fragmentation (Fig. 3I-3J) (*28*). Mitochondria also often appeared concentrated in the perinuclear region, and, as in the case of MYO19 depletion, mitochondria failed to traffic to filopodia (Fig. 3K). Despite the pleiotropic effects of MIC60 depletion, we observed that while filopodia density was not significantly altered, filopodia length was reduced (Fig. 3L-3M), consistent with our observations in MYO19-depleted cells. We also immunoblotted and immunolabeled MIC60-depleted cells for MYO19, finding that MYO19 levels were minimally affected and MYO19 retained its targeting to the OMM (Fig. S2J-S2K). Together, these data indicate that disruption of the MICOS complex or depletion of MYO19 each interfere with METEOR formation and mitochondrial trafficking to filopodia. Further, in both cases, the loss of mitochondrial targeting to filopodia correlates with a reduction in filopodia length.

### METEOR mitochondria have a unique composition

The MICOS complex is typically enriched within the IMM at cristae junctions, explaining the semi-punctate distribution of its subunits in immunofluorescence images (*33*). Based on this, we predicted that the local enrichment of MICOS and MIB subunits within METEORs would reflect an enrichment of cristae membranes, and thus respiratory complexes, at those sites. To assess the local distribution of respiratory complex proteins at MICOS enrichments, we fixed and immunolabeled wild-type cells for MIC60 or MIC19 and constituent members of Complex I (NDUFB8), Complex III (UQCRFS1), Complex IV (COX4I1), and ATP synthase (ATP5F1A). We then measured the fluorescence intensity ratio of each respiratory complex component at MICOS enrichment sites relative to nearby mitochondria in the cell periphery. To our surprise, respiratory complex subunit staining was not enriched at METEORs versus other mitochondria and did not appear significantly different than the IMM protein TIMM23 (Fig. 4A-4B). These data suggest that the local enrichment of MICOS subunits does not reflect an enrichment of respiratory complexes within METEORs.

**Figure 4.**
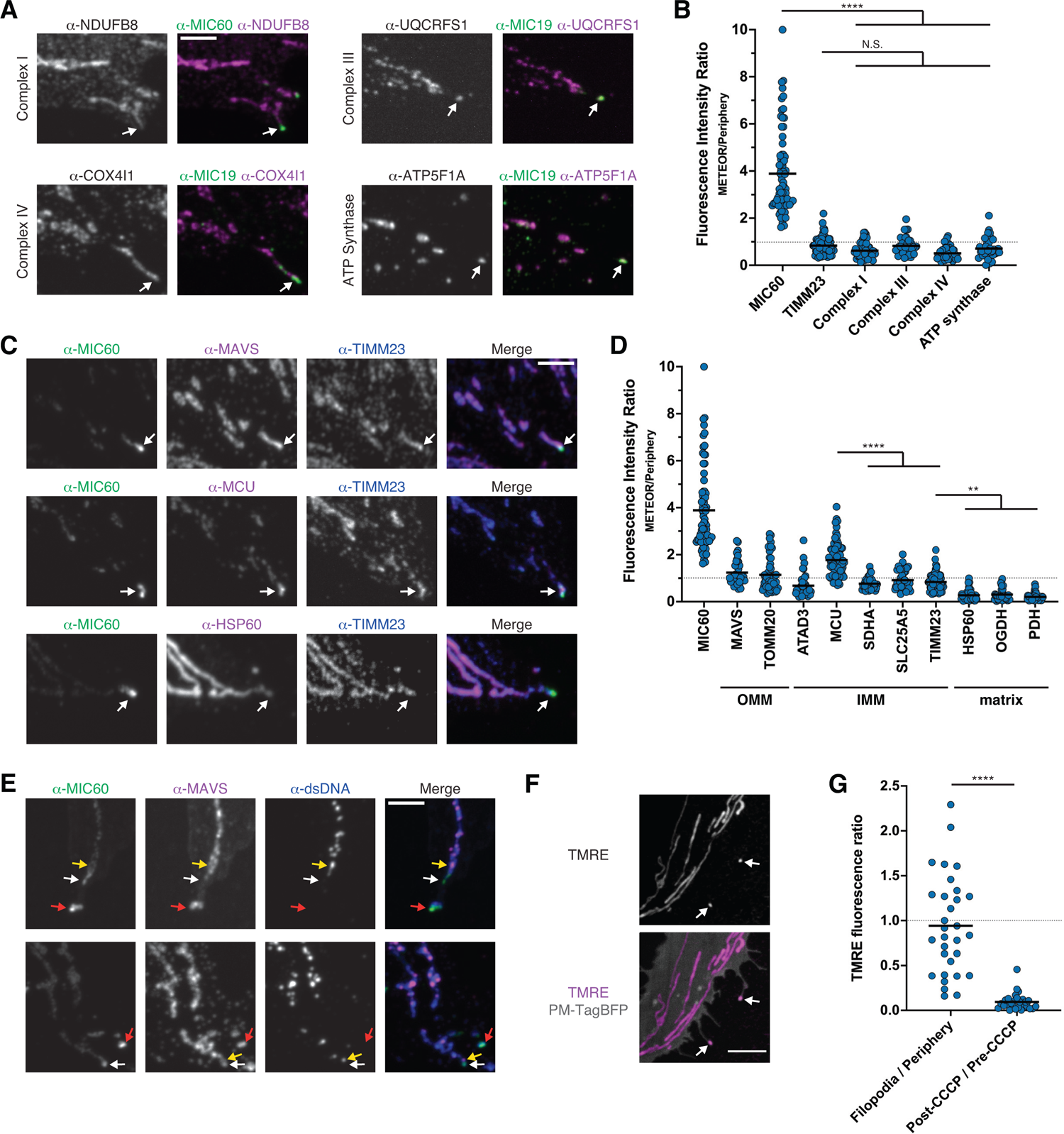
METEOR mitochondria have a unique composition. **(A)** Maximum intensity projection confocal images are shown of fixed U2OS cells immunolabeled for MIC60 or MIC19 and the indicated respiratory complex subunit. White arrows mark sites of MICOS enrichment. **(B)** A graph of fluorescence intensity ratios at METEOR tips relative to nearby cell peripheral mitochondria for MIC60 (n=71), TIMM23 (n=76), Complex I (NDUFB8; n=56), Complex III (UQCRFS1; n=38), Complex IV (COX4I1; n=47), and ATP Synthase (ATP5F1A; n=35). Data for MIC60 and TIMM23 are redisplayed from Fig. 1E. Images were acquired from at least two independent experiments. Solid lines indicate means and a dotted line is shown at y=1 for reference. **(C)** Maximum intensity projection images are shown of cells immunolabeled for MIC60, TIMM23, and either MAVS, MCU, or HSP60, where indicated. White arrows mark sites of MIC60 enrichment. Note the apparent uniform labeling of MAVS (top panel), co-enrichment of MCU (middle panel), and de-enrichment of HSP60 (bottom panel). **(D)** A graph as in (B) for MIC60 (n=71), MAVS (n=38), TOMM20 (n=49), ATAD3 (n=41), MCU (n=82), SDHA (n=38), SLC25A5 (n=38), TIMM23 (n=76), HSP60 (n=44), OGDH (n=43), and PDH (n=44). Data for MIC60, TIMM23, and TOMM20 are redisplayed from Fig. 1E. Images were acquired from at least two independent experiments. Solid lines indicate means and a dotted line is shown at y=1 for reference. **(E)** As in (C) for cells immunolabeled for MIC60, MAVS, and dsDNA. White arrows mark MIC60 enrichment sites within mitochondria that contain mtDNA (yellow arrows), but mtDNA foci are not localized at those sites. Red arrows depict METEOR mitochondrial fragments that do not contain mtDNA. **(F)** A single plane live cell confocal image of a U2OS cell transfected with PM-TagBFP and labeled with TMRE. The white arrows indicate mitochondria in filopodia. **(G)** A graph depicting the relative fluorescence intensity ratios of TMRE labeling in mitochondria in filopodia relative to cell peripheral mitochondria for cells treated as in (F) (n=31 from 17 cells). For comparison, the fluorescence intensity ratio was determined for mitochondria in the cell periphery that were imaged before and after 45 minutes of treatment with 10 µM CCCP (n=27 from 20 cells). Solid lines indicate means. A dotted line is shown at y=1 for reference. Asterisks (****p<0.0001, **p<0.01) represent ordinary one-way ANOVAs with Dunnett’s multiple comparisons test (B, D) or an unpaired two-tailed *t* test (G). N.S. indicates not statistically significant. Scale bars: (A,C,E) 3 µm; (F) 5 µm. See also Figure S3.

Given the unusual distribution of MICOS in METEORs and the surprising lack of co-enrichment of respiratory complex subunits at these sites, we evaluated the protein composition of METEORs in more detail. We took a candidate-based approach by utilizing antibodies against proteins in different mitochondrial sub-compartments that clearly labeled the organelle by immunofluorescence. Aside from MYO19 and the MIB subunits SAMM50 and DNAJC11 (Figs. 3 and S1), other OMM proteins such as TOMM20 and MAVS were not enriched at METEOR tips (Fig. 4C-4D). In contrast, the IMM and matrix proteins we examined often exhibited either selective enrichment or depletion. Specifically, each matrix-localized protein we assessed appeared de-enriched as compared to IMM proteins such as TIMM23. These included the chaperone HSP60, the citric acid cycle enzyme α-ketoglutarate dehydrogenase (OGHD), and the pyruvate dehydrogenase E1α subunit (PDHA1) (Fig. 4C-4D; Fig. S3). We also saw differential labeling of IMM proteins. Nearly every IMM component we analyzed had a similar staining pattern as TIMM23, including the ATP-ADP transporter SLC25A5, the AAA ATPase ATAD3, and the peripherally associated succinate dehydrogenase subunit SDHA (Fig. 4D; Fig. S3). In contrast, we saw an almost 2-fold enrichment of the Ca^2+^ uniporter MCU at METEOR tips (Fig. 4C-4D). Indeed, MCU was the only other protein we examined that exhibited substantial local enrichment within METEORs. In total, our data suggest that METEORs have a distinct composition from other mitochondria in the cell.

Because of the general reduction in relative protein staining of matrix-localized subunits, we assayed for the presence of mtDNA at METEOR tips. We immunolabeled cells with dsDNA antibody, which labels both nuclear DNA as well as mitochondrial genomes. Notably, while mtDNA always appeared present in mitochondrial tubules where MICOS was enriched at the tip, the mtDNA never localized at the enrichment site (0%, n=60; Fig. 4E, compare yellow and white arrows). Further, METEORs frequently appear as mitochondrial fragments, and in these cases, mtDNA was infrequently present and also never localized to a MIC60 enrichment site (Fig. 4E, see red arrows). Thus, in addition to having relatively low abundance of matrix proteins, METEOR tips and METEOR fragments generally do not contain mtDNA.

Due to the unusual composition of METEOR tips, their relatively low levels of matrix proteins, and the absence of mtDNA, we considered that these mitochondria may not be functional. Indeed, previous reports suggest that in neutrophils, MYO19 mediates trafficking of dysfunctional mitochondria with low membrane potential to migrasomes for their disposal, a process termed mitocytosis (*52*). We therefore examined the membrane potential of mitochondria in filopodia, which are almost exclusively METEORs (Fig. 2G). We transfected cells with PM-TagBFP and stained mitochondria with the membrane potential-dependent dye tetramethylrhodamine ethyl ester (TMRE). We observed equivalent TMRE staining in mitochondria in filopodia relative to cell peripheral mitochondria (Fig. 4F-4G). By comparison, treatment with carbonyl cyanide m-chlorophenyl hydrazone (CCCP), which uniformly collapses membrane potential, ablated TMRE signal (Fig. 4F-4G). These data suggest that despite their unusual composition, METEOR mitochondria in filopodia maintain polarization across the IMM.

### Elimination of mitochondria from filopodia disrupts cellular migration

Filopodia are functionally associated with cell adhesion and migration (*41, 42*). Given the correlation between mitochondrial presence and filopodia length, we considered that the compositionally unique mitochondria could promote filopodia function and support cell migration. To examine this, we imaged cells over a period of 3.5 hours. Nuclei stained with Hoechst were used to determine the individual motility of several hundred cells per experiment. When plated on uncoated glass-bottom dishes, a small percentage of cells was motile as determined by their displacement rate, whereas cellular movement on a population level was relatively limited (Fig. 5A-5B). However, consistent with previous reports (*53*), when plated on a surface pre-coated with collagen, U2OS cell migration was substantially increased, and individual cellular motility was more uniform over the population of cells (Fig. 5A-5B and Video 6).

**Figure 5.**
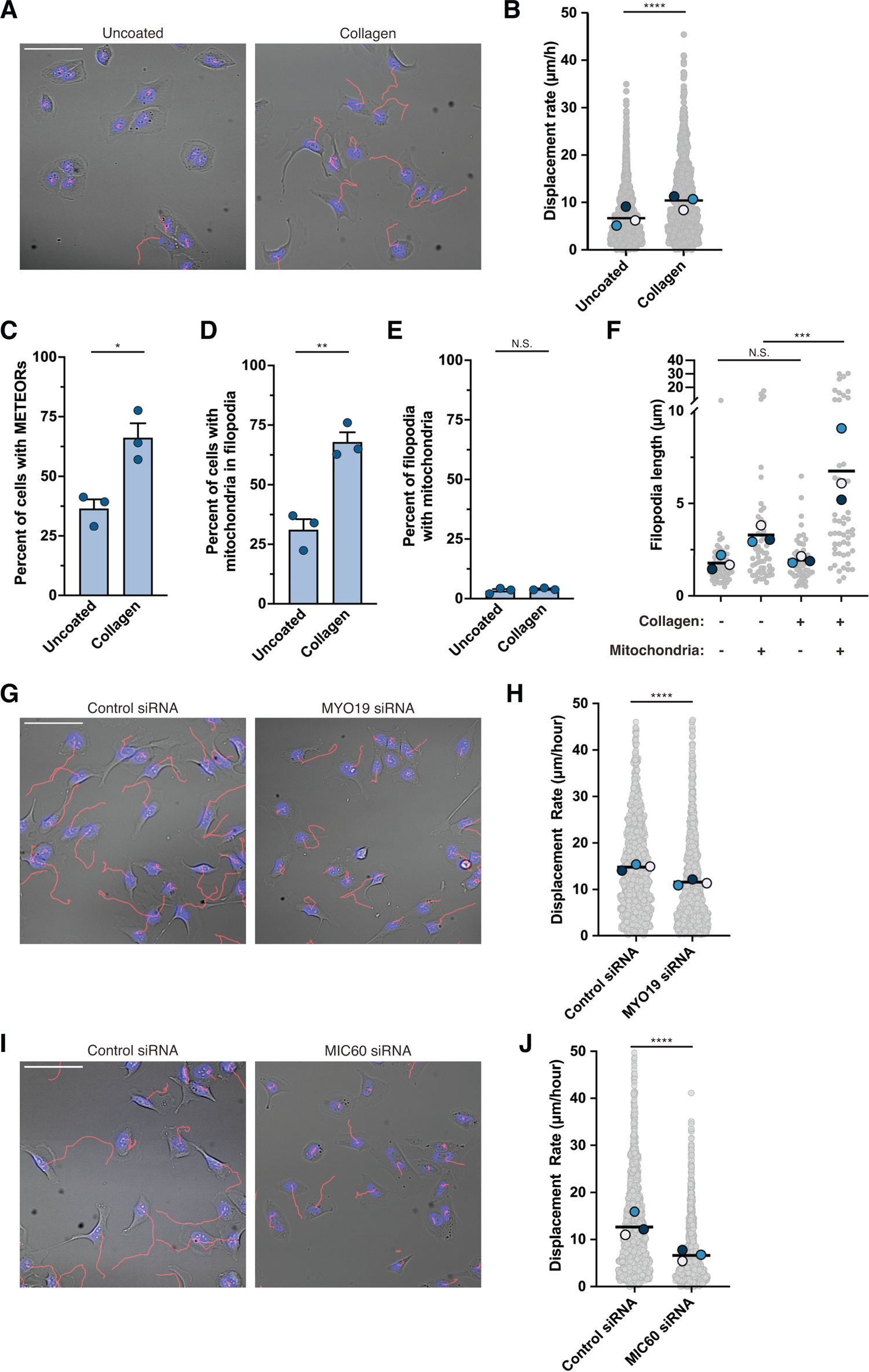
Elimination of mitochondria from filopodia disrupts cellular migration. **(A)** Representative images are shown of U2OS cells plated on uncoated (left) or collagen coated (right) glass-bottom dishes, labeled with Hoechst (blue), and imaged every 5 minutes. Red lines indicate nuclear movement over a period of 210 minutes. See Video 6. **(B)** A graph depicting the displacement rate of individual cells as in (A). Data represent at least ∼1000 cells per condition acquired from three independent experiments. Solid lines indicate mean displacement rate and superimposed colored circles represent the means of individual experimental replicates. **(C-E)** Quantitative analyses are shown of cells that were plated on uncoated or collagen coated dishes, fixed, stained with fluorescence-conjugated WGA, immunolabeled with MIC60 and TIMM23, and imaged by confocal microscopy. Graphs are shown depicting (C) the frequency of cells with METEORs, (D) the frequency of cells with mitochondria in filopodia, and (E) the frequency of filopodia that contain mitochondria. Data shown represent ∼50 cells per condition in each of three independent experiments and bars indicate S.E.M. **(F)** A graph depicting the individual lengths of filopodia that contain mitochondria, or adjacent filopodia without mitochondria, from cells as imaged in (C-E). Data represent ∼50 filopodia per condition. Solid lines indicate the mean filopodia length. Superimposed colored circles represent the means for individual experimental replicates. **(G)** As in (A) for cells treated with control siRNA (left) or siRNA targeting MYO19 (right). See Video 7. **(H)** A graph depicting cellular displacement rate as in (B) for cells treated as in (G). **(I-J)** As in (G-H) for cells treated with control siRNA (left) or siRNA targeting MIC60 (right). See Video 8. Asterisks (****p<0.0001, ***p<0.001, **p<0.01, *p<0.05) represent unpaired two-tailed *t* tests (B,C,D,E,H,J) or an ordinary one-way ANOVA with Tukey’s multiple comparisons (F). N.S. indicates not statistically significant. Scale bars: (A, G, I) 100 µm.

Given the increased migration of U2OS cells plated on a collagen substrate, we asked whether the increased rate of motility positively correlates with the presence of METEORs. We grew wild-type cells on uncoated or collagen-coated glass-bottom dishes, stained the PM post-fixation, and immunolabeled mitochondria with TIMM23 and MIC60. Strikingly, while only about one-third of cells exhibited METEORs when plated on a glass substrate, they were present in approximately two-thirds of cells plated on collagen (Fig. 5C). This corresponded to a similar increase in the percentage of cells with mitochondria present in filopodia, but notably, mitochondria remained present in only a minor subset of filopodia in each cell (Fig. 5D-5E). Based on our observation that filopodia with mitochondria present were longer than other filopodia (Fig. 2H), we next asked whether plating cells on collagen affected filopodia length. We found that the length of filopodia that do not contain mitochondria, which comprise ∼95% of the population, was not affected by plating cells on collagen (1.9 µm collagen-coated vs. 1.8 µm uncoated; Fig. 5F).

However, remarkably, the mean length of filopodia that contain mitochondria was approximately twice as long after plating cells on collagen (6.8 µm collagen-coated vs. 3.3 µm uncoated; Fig. 5F). Together, these data indicate that plating cells on a substrate that promotes cell migration increases the prevalence of METEORs in filopodia and correlates with a selective increase in length of filopodia that contain mitochondria.

Since the presence of METEORs positively correlated with cell migration rates, we next determined whether inhibition of the formation of METEORs and elimination of mitochondria from filopodia affected cellular migration. We knocked down MYO19, plated cells on dishes coated with collagen, and analyzed cellular migration as before. Depletion of MYO19 led to a significant reduction in the cellular displacement rate, indicating that MYO19 is required for normal cellular migration (Fig. 5G-5H, Video 7, and Fig. S2L-S2M). However, because MYO19 knockout cells are reported to have defects in focal adhesions (*49*), we considered that MYO19 knockdown may influence cell migration independently of METEOR formation. We thus reasoned that if the reduced migration rate of MYO19-depleted cells was due to the inability of METEORs to traffic to filopodia, then knockdown of MIC60, which also prevents mitochondrial targeting to filopodia without affecting MYO19 recruitment (Fig. 3K and Fig. S2K), would decrease cell motility. We performed siRNA-mediated knockdown of MIC60 and examined the rate of migration of cells grown on dishes coated with collagen. As in the case of MYO19-depleted cells, disruption of the MICOS complex via MIC60 knockdown led to a significant reduction of cellular migration (Fig. 5I-5J and Video 8). Reduced migration in MIC60-depeleted cells may be due to pleiotropic and global defects in mitochondrial function. However, in combination with our observations of reduced cellular migration in MYO19-depleted cells, these data are consistent with the model that filopodia targeting of METEORs is required to promote normal cellular motility rates.

## Discussion

Within a single cell, mitochondria are heterogeneous in size, shape, protein composition, cristae organization, and inter-organelle contacts. Here, we identify a subpopulation of compositionally unique mitochondria, METEORs, that localize in proximity to the PM and within filopodia. These mitochondria are dramatically remodeled. Their tips are highly enriched for the MIB/MICOS complex and the IMM Ca^2+^ uniporter MCU while also having reduced levels of matrix proteins and mitochondrial genomes. We find that these mitochondria traffic to filopodia in a manner dependent on MYO19, which co-enriches with the MICOS complex at METEORs. Further, we observe a positive correlation between the length of filopodia and the presence of mitochondria within them. Growth on a collagen substrate promotes mitochondrial trafficking to longer filopodia while depletion of MYO19 or disruption of MICOS prevents mitochondrial trafficking to PM extensions, resulting in a selective reduction of longer filopodia. Finally, we find that elimination of mitochondria from filopodia by either approach is associated with reduced rates of cellular motility. Together, our data indicate that a specialized sub-population of mitochondria in the cell periphery localizes to filopodia to support cellular migration.

Cell motility is a fundamentally vital process, essential for tissue and organ development and wound healing, and is often dysregulated in metastatic cancer cells (*54*). Mitochondria are known to traffic to the leading edge and have been positively implicated in supporting migration and cell invasion via energy production (*55–58*). In addition, previous reports suggest a role for MYO19 in trafficking of mitochondria to filopodia and increasing filopodia formation during stress conditions (*44, 45*). Our findings suggest that mitochondria can target to filopodia not only during stress, but also during basal growth conditions to support migration. It is thought that mitochondria traffic to the leading edge to increase the local concentration of ATP required for energy demanding processes. However, we show that the mitochondria that are found within filopodia are highly unusual. High levels of MICOS complexes canonically would predict increased cristae density, which would correspond to increased respiratory complex concentration and higher OXPHOS function. Notably, we do not observe respiratory complex enrichment nor the presence of mtDNA at MICOS-enriched sites. Importantly, METEORs do generally maintain membrane potential, suggesting they are indeed functional. Interestingly, out of the 14 mitochondrial proteins we examined, the only protein we regularly observed co-enriched with MICOS/MIB/MYO19 within METEORs was the IMM Ca^2+^ uniporter MCU. In neuronal development, cytosolic Ca^2+^ transients within filopodia are important for regulating growth cone motility and guidance (*59*) and other reports suggest that PM L-type Ca^2+^ channels promote a local influx of Ca^2+^ within filopodia that promotes their stability and cell migration (*60, 61*). An intriguing possibility is that MCU-remodeled mitochondria localize within filopodia to help facilitate regulation of migration, either by utilizing Ca^2+^ to support OXPHOS or by playing an energy production-independent Ca^2+^ homeostasis role (*62, 63*). While we have observed mitochondrial localization to filopodia in U2OS cells under standard growth conditions, it remains an open question whether METEOR formation occurs in other cell types or processes, for example during neuronal development.

A critical question that arises from our work is the mechanism of protein remodeling and MICOS enrichment at METEORs. Our results suggest that depletion of MYO19 prevents not only mitochondrial trafficking to filopodia, but also METEOR formation. Thus, the remodeling of the mitochondrial proteins may be initiated from the outside-in. One interesting possibility that warrants further exploration is whether MICOS influences the redistribution of other IMM/matrix proteins that occurs within METEORs. While the MICOS complex is most associated with the formation and stabilization of cristae junctions, MICOS has a broader distribution within the IMM than merely at cristae junctions (*34*). MICOS also serves as a protein-protein interaction hub within the intermembrane space and has previously been shown to interact with the MCU regulatory subunit MICU1 (*23, 29*). Thus, it will be interesting to determine whether via this interaction, MICOS is specifically required for the local enrichment of MCU. Another question that arises is how matrix proteins and mtDNA are more generally excluded from METEOR tips. Going forward, it will be pivotal to determine if the localization of MICOS in METEORs can be uncoupled from the protein remodeling that occurs there, which will improve our understanding of the specific functional roles of METEORs.

Our work also has important implications for our understanding of the subcellular heterogeneity of mitochondria. In recent years, it has become clear that mitochondria are specialized and proteomically or functionally distinct in different cell types and tissues (*8, 9*). Mitochondria also can have different morphological and protein density characteristics at a subcellular level (*7, 33*). There are also examples of remodeled mitochondrial membranes, including MDVs, MDCs, and SPOTs, that can serve in signaling and stress responses (*38*). Among our findings, we notably identify a novel subtype of mitochondria within a single cell with a distinct composition, adding to the growing list of specialized mitochondrial functions and behaviors. Our results stem from a fortuitous observation by immunofluorescence microscopy of local enrichment of a single protein at rare sites within the mitochondrial network. This raises the intriguing possibility that such examples of organelle specialization are more widespread throughout nature than previously thought and may only become apparent by careful unbiased microscopy-based approaches.

## Supplemental Data

**Figure S1.**
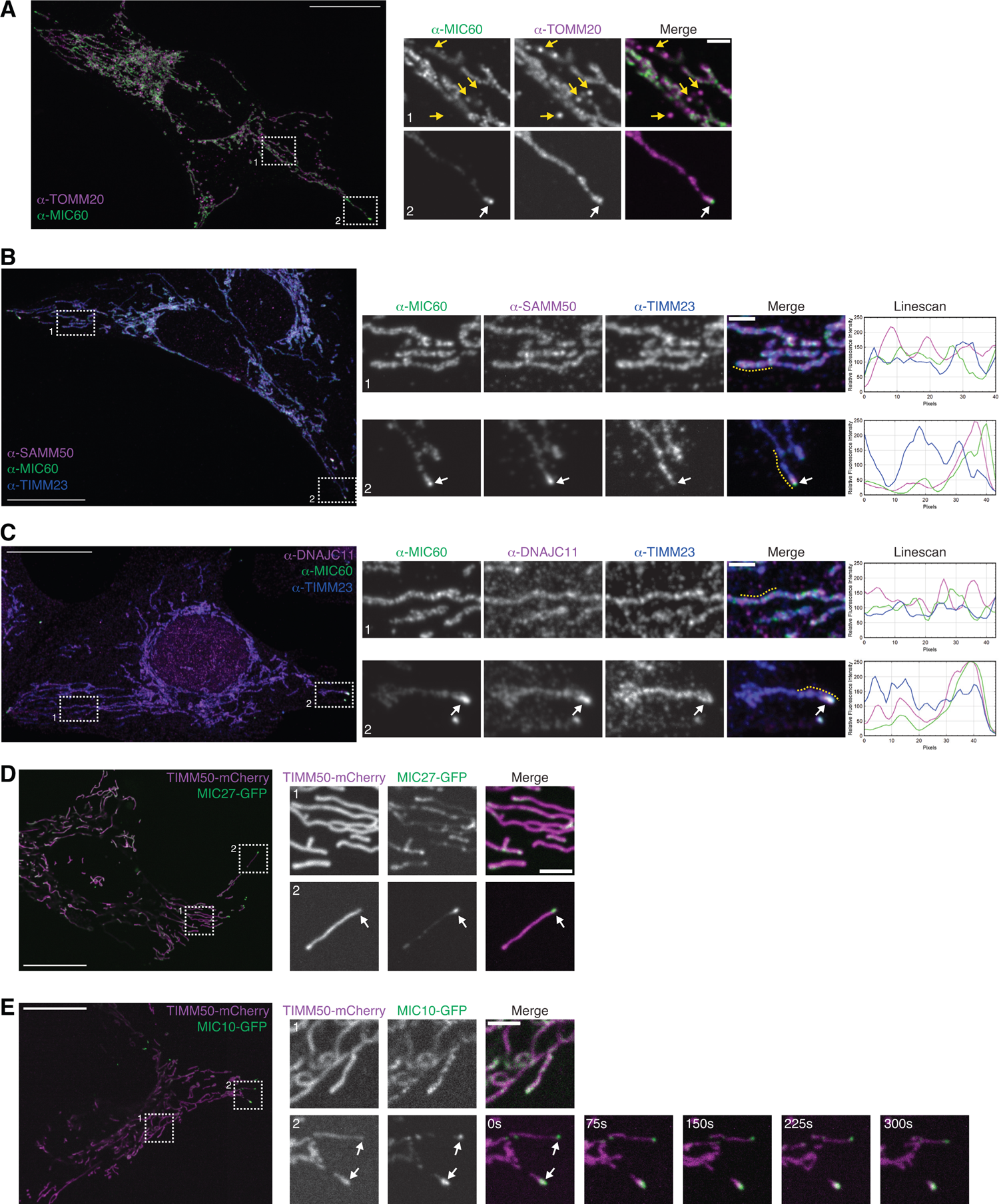
MICOS and MIB subunits are enriched at the tips of METEORs. **(A)** Maximum intensity projection confocal images are shown of a fixed U2OS cell immunolabeled for MIC60 and TOMM20. Dotted boxes correspond to the numbered insets shown on the right. Box 1 shows a region in the perinuclear area of the cell and Box 2 shows a METEOR. White arrows mark a site of MIC60 enrichment and yellow arrows mark MDVs. **(B-C)** Images are shown as in (A) for cells immunolabeled for MIC60, TIMM23, and SAMM50 (B) or DNAJC11 (C). Box 1 shows a representative mitochondrion in the cell periphery and Box 2 shows a METEOR. The yellow dotted lines on the insets correspond to the fluorescence intensity linescans shown at the right. White arrows mark sites of MICOS enrichment. **(D)** Single-plane confocal images are shown of a U2OS cell transiently transfected with low-levels of TIMM50-mCherry and MIC27-GFP. Dotted boxes correspond to the numbered insets shown on the right. Box 1 shows representative mitochondria in the cell periphery and Box 2 shows a MIC27-enriched METEOR. Box 2 images correspond to those shown in Fig. 1F and Video 2. **(E)** As in (D) for a cell transfected with TIMM50-mCherry and MIC10-GFP. At right, timelapse microscopy images are shown of Box 2 at the indicated timepoints (s = seconds). Scale bars: (A,B,C) 20 µm; (2 µm in inset); (D,E) 20 µm (3 µm in inset).

**Figure S2.**
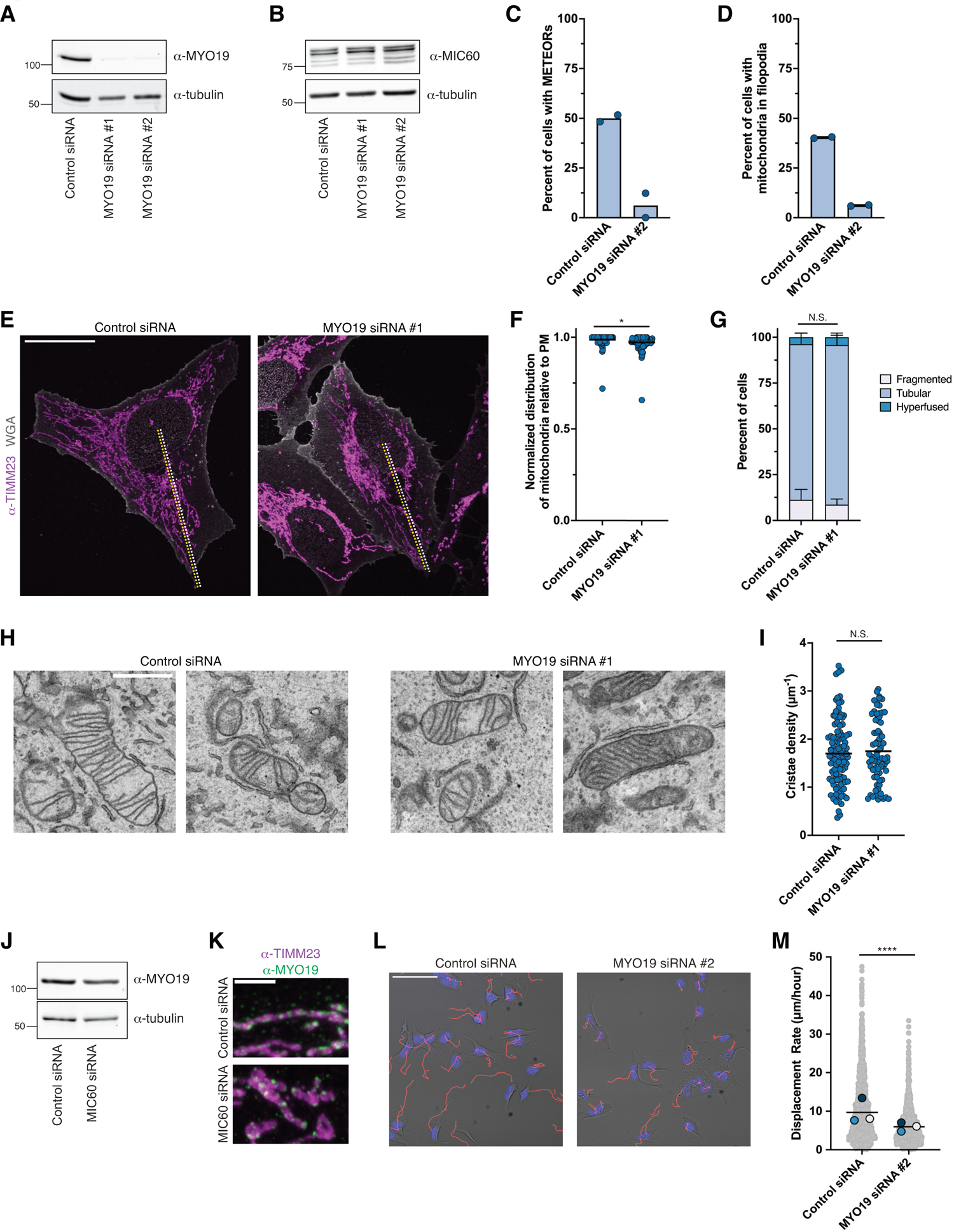
MYO19 depletion does not grossly affect mitochondrial distribution or cristae morphology. **(A-B)** Western blots analyses with the indicated antibodies of lysate from U2OS cells transiently transfected with control siRNA or the indicated siRNAs targeting MYO19. **(C-D)** A graph depicting the frequency of cells with METEORs (C) or mitochondria in filopodia (D) from cells treated as in (A-B) with MYO19 siRNA #2. Cells were fixed and immunolabeled with TIMM23, MIC60, and fluorescence-conjugated WGA as in Fig. 3D. Data shown represent >30 cells per condition in each of two independent experiments. Experiments were performed simultaneously with those in Fig. 3E-3F and control siRNA data are redisplayed. **(E-F)** Maximum intensity projection confocal images are shown (E) of fixed cells treated as in Fig. 3C with MYO19 siRNA #1 and immunolabeled for TIMM23 and stained with fluorescence-conjugated WGA. A graph is shown in (F) depicting the ratio of the distance from mid-nucleus to furthest cell peripheral mitochondrion (see yellow lines in (E)) relative to the distance between mid-nucleus and the PM (see white lines in (E)) for individual cells. Data shown represent ∼80 total cells per condition acquired from three independent experiments. Solid lines represent means. **(G)** A graph depicting the frequency of cells with the indicated mitochondrial network character categorized from cells treated as in Fig. 3C and imaged as in (E). Data shown represent ∼30-60 cells per condition in each of three independent experiments and bars indicate S.E.M. **(H-I)** Representative electron microscopy images of mitochondria are shown (H) of cells treated with control siRNA or MYO19 siRNA #1 as in Fig. 3C. A graph is shown (I) depicting the cristae density (number of cristae per µm of mitochondrial perimeter) from images as in (H). Data shown represent individual mitochondrial profiles (n=109 control and 77 MYO19 siRNA-treated mitochondria from approximately 8 cells per condition). Solid lines represent means. **(J)** Western blot analysis with the indicated antibodies of lysate from cells transfected with the indicated siRNAs. **(K)** Maximum intensity projection confocal images are shown of cells treated as in (J) and fixed and immunolabeled with TIMM23 and MYO19. **(L)** Representative images are shown of cells treated as in (A) with MYO19 siRNA #2, plated on collagen coated glass-bottom dishes, labeled with Hoechst (blue), and imaged every 5 minutes. Red lines indicate nuclear movement over a time-course of 210 minutes. **(M)** A graph depicting the displacement rate of individual cells as in (L). Data represent at least ∼1000 cells per condition acquired from three independent experiments. Solid lines indicate mean displacement rate and superimposed colored circles represent the means of individual experimental replicates. Asterisks (****p<0.0001, *p<0.05) represent unpaired two two-tailed *t* tests (F, I, M) or a two-way ANOVA with Šídák’s multiple comparisons test (G). N.S. indicates not statistically significant. Scale bars: (E) 25 µm; (H) 1 µm; (K) 3 µm; (L) 100 µm.

**Figure S3.**
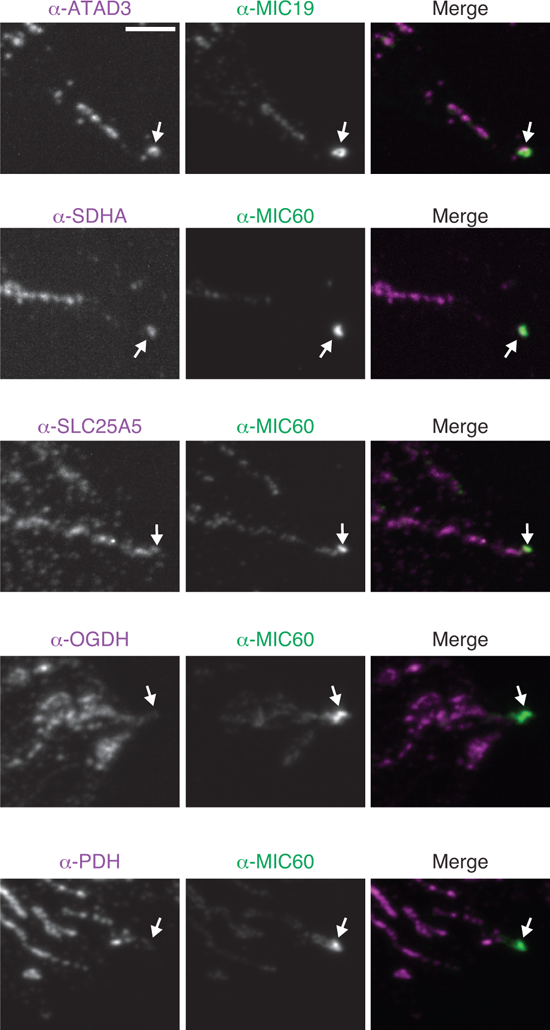
METEORs have a unique protein composition. Maximum intensity projection confocal images are shown of fixed U2OS cells immunolabeled with the indicated antibodies. White arrows mark sites of MICOS enrichment. Images correspond to quantifications shown in Figure 4D. Scale bar: 3 µm.

**Video 1. Timelapse microscopy of stable MIC27 enrichment at the tip of a dynamic mitochondrion.** A video of timelapse microscopy images of a mitochondrion transiently transfected with TIMM50-mCherry (magenta) and MIC27-GFP (green). Images are single confocal planes captured at the indicated time intervals. Still images of Video 1 are shown in Fig. 1F (top panel). Scale bar: 3 µm.

**Video 2. Timelapse microscopy of dynamic MIC27 redistribution at the tip of a METEOR.** A video of timelapse microscopy images of a cell transiently transfected with TIMM50-mCherry (magenta) and MIC27-GFP (green). Images are single confocal planes captured at the indicated time intervals. Still images are shown in Fig. 1F (middle panel) and Fig. S1D. Scale bar: 3 µm.

**Video 3. Timelapse microscopy of a MIC27-labeled METEOR that undergoes fission.** A video of timelapse microscopy images of a cell transiently transfected with TIMM50-mCherry (magenta) and MIC27-GFP (green). Images are single confocal planes captured at the indicated time intervals. Still images are shown in Fig. 1F (bottom panel). Scale bar: 3 µm.

**Video 4. Timelapse microscopy of a METEOR that dynamically localizes to a growing plasma membrane extension.** A montage of timelapse microscopy images of a cell transiently expressing TIMM50-mCherry (magenta), MIC27-GFP (green), and PM-TagBFP (gray). Images shown are single confocal planes captured at the indicated time intervals. Still images are shown in Fig. 2C (top panel). Scale bar: 3 µm.

**Video 5. Timelapse microscopy of a METEOR that dynamically moves away from the tip of a plasma membrane extension.** A montage of timelapse microscopy images of a cell transiently expressing TIMM50-mCherry (magenta), MIC27-GFP (green), and PM-TagBFP (gray). Images shown are single confocal planes captured at the indicated time intervals. Still images are shown in Fig. 2C (bottom panel). Scale bar: 3 µm.

**Video 6. A comparison of motility of cells plated on uncoated or collagen-coated microscopy dishes.** A montage of timelapse brightfield microscopy images of U2OS cells grown on uncoated dishes (left) or collagen-coated dishes (right) at the indicated time intervals. The video corresponds to images shown in Fig. 5A. Scale bar: 100 µm.

**Video 7. A comparison of motility of control siRNA treated cells or cells depleted of MYO19.** A montage of timelapse brightfield microscopy images of U2OS cells treated with control siRNA (left) or MYO19-targeting siRNA (right) at the indicated time intervals. The video corresponds to images shown in Fig. 5G. Scale bar: 100 µm.

**Video 8. A comparison of motility of control siRNA treated cells or cells depleted of MIC60.** A montage of timelapse brightfield microscopy images of U2OS cells treated with control siRNA (left) or MIC60-targeting siRNA (right) at the indicated time intervals. The video corresponds to images shown in Fig. 5I. Scale bar: 100 µm.

## Materials and Methods

### Cell culture

U2OS cells (a gift from Jodi Nunnari) were originally acquired from ATCC and were cultured in DMEM (D5796; Sigma Aldrich) supplemented with 10% fetal bovine serum (F0926; SigmaAldrich), 25 mM HEPES (H0887; SigmaAldrich), 100 units/mL penicillin, and 100 µg/mL streptomycin. Cells were maintained at low passage number and routinely tested for mycoplasma contamination.

### Plasmids and siRNA oligonucleotides

MIC27-GFP was generated by PCR amplifying the MIC27 coding sequence (NCBI Accession Number NM_198450.6) from cDNA generated from HeLa cells and cloning into the BglII/SalI sites of pAcGFP-N1. MIC27-mCherry was generated by inserting mCherry into the KpnI/NotI sites of MIC27-GFP, replacing the GFP cassette. MIC10-GFP was generated by PCR amplifying the MIC10 isoform a coding sequence (NCBI Accession Number NM_001032363.4) from cDNA generated from HeLa cells and cloning into the XhoI/BamHI sites of pAcGFP-N1. TIMM50-AcGFP was previously described (*39*). PM-TagBFP was generated by first PCR amplifying an N-terminal fusion of the PM targeting sequence of Lyn (*40*) to the AcGFP coding sequence from pAcGFP-N1 to generate PM-GFP. To generate PM-TagBFP, TagBFP was subsequently subcloned into the BamHI/NotI sites of PM-GFP, replacing the GFP cassette. Myo19-GFP (Addgene 134988) was a gift from Martin Bähler (*64*). GFP-Myo10 (Addgene 135403) was a gift from Richard Cheney (*43*).

For siRNA depletion, Silencer Select siRNAs (ThermoFisher Scientific; Negative Control No. 2 (4390846); MIC60 (s21633); MYO19 (siRNA#1 - s37017; siRNA#2 - s37019)) were used for all experiments. siRNA sequences are as follows: MIC60: 5’-GAAUGACCUAGAAACGAAtt-3’, MYO19 #1: 5’-GGUGAAUCCUGUGACACUAtt-3’, MYO19 #2: 5’-GCGUGUACACUGAGGAAUAtt-3’

### Transient transfections and siRNA treatments

For transient expression of plasmids, approximately 150,000 cells were seeded per well of a 6-well dish and allowed to adhere overnight prior to transfection. The next day, 30-100 ng of plasmid DNA was transfected using Lipofectamine 3000 (ThermoFisher Scientific) according to the manufacturer’s instructions for 5-8 hours. Cells were then passaged to microscope dishes and allowed to adhere overnight prior to imaging.

For siRNA treatments, approximately 150,000 cells were seeded per well of a 6-well dish and allowed to adhere overnight prior to transfection. The next day, Lipofectamine RNAiMAX (ThermoFisher Scientific) and the indicated siRNA oligonucleotides (20 nM) were applied to the cells according to the manufacturer’s instructions. Cells were incubated for 24 hours, dissociated with trypsin, diluted 1:2 to new dishes, allowed to recover for 24 hours, and re-transfected for 5 hours with RNAiMAX and siRNA oligonucleotides (20nM). Cells were then passaged to glass-bottom microscope dishes or standard growth dishes and incubated ∼24 hours before imaging or Western analysis.

### Whole-cell lysates and Western analysis

To prepare whole-cell lysates, cells were dissociated with trypsin, washed once with DPBS, and lysed in 1X RIPA buffer (150mM NaCl, 50mM Tris HCl pH 7.5, 1% sodium deoxycholate, 0.1% SDS, 1% NP-40, 1mM EDTA) supplemented with 1X protease-inhibitor cocktail (539131; MilliporeSigma). Protein concentration was determined using a BCA assay (BioRad) and normalized between samples prior to adding 6X Laemmli buffer [6% SDS (w/v). 21.6% glycerol (v/v), 0.18M Tris HCl, pH 6.8, 0.01% bromophenol blue (w/v), and 10% β-mercaptoethanol (v/v)] to a final concentration of 1X. Samples were heated for 5 minutes at 95°C and lysate was resolved on Tris-Glycine polyacrylamide gels. After electrophoresis, proteins were transferred to PVDF membranes (0.45µm pore size) and immunoblotted with the following primary antibodies (mouse anti-MIC60 [110329; Abcam], rabbit anti-MYO19 [ab174286; Abcam], mouse anti-Tubulin [66031-1-Ig; ProteinTech]). The appropriate secondary antibodies (anti-rabbit DyLight800 [SA5-35571; ThermoFisher Scientific], or goat anti-mouse DyLight680 [35568; ThermoFisher Scientific]) were used and images were acquired with a ChemiDoc MP Imaging System (BioRad).

### Immunofluorescence staining

For all immunofluorescence assays, ∼50,000 cells were plated on glass-bottom cover dishes (CellVis D35-14-1.5-N). Where indicated, glass-bottom dishes were first coated with 1 mL of 0.05 mg/mL Bovine Collagen I (354231; Corning) for 1 hour. Collagen was aspirated and dishes were allowed to dry for 15-20 minutes. Adherent cells were fixed in 4% paraformaldehyde in PBS (15 min, room temperature), except for ATP5F1A labeling, where cells were fixed with ice cold methanol for 15 minutes. Where indicated, cells were incubated with a 1:100 dilution of WGA-CF405s (290271; Biotium) in Hank’s Balanced Salt Solution (SigmaAldrich) for 30 minutes to label the PM. Cells were then permeabilized (0.1% Triton X-100 in PBS) for 5 minutes and incubated in blocking solution (10% FBS and 0.1% Triton X-100 in PBS) for 30 minutes. Cells were then incubated for 30 minutes at room temperature or overnight at 4°C with the indicated primary antibodies diluted 1:100-1:200 in blocking solution (mouse IgG1 anti-MIC60 [110329; Abcam], mouse IgG2a anti-TIMM23 [611222; BD Biosciences], mouse IgG2α anti-TOMM20 [sc17764; Santa Cruz Biotechnology], rabbit anti-APOOL (MIC27) [PA5-51427; ThermoFisher Scientific], rabbit anti-CHCHD3 (MIC19) [HPA042935; Atlas Antibodies], rabbit anti-DNAJC11 [ab183518; Abcam], rabbit anti-SAMM50 [HPA034537; Atlas Antibodies], rabbit anti-SLC25A5 [HPA046835; Atlas Antibodies], rabbit anti-NDUFB8 [14794-1-AP; Proteintech], mouse IgG1 anti-UQCRFS1 [sc-271609; Santa Cruz Biotechnology], mouse IgG1 anti-COX4I1 [66110-1-Ig; Proteintech], rabbit anti-ATP5F1A [ab128743; Abcam], rabbit anti-PDH E1 Alpha [18068-1-AP; Proteintech], rabbit anti-OGDH [15212-1-AP; Proteintech], rabbit anti-HSP60 [15282-1-AP; Proteintech], rabbit anti-MCU [14997; Cell Signaling Technology], mouse IgG2α anti-dsDNA [ab27156; Abcam], rabbit anti-MYO19 [ab174286; Abcam], rabbit anti-MAVS [24930; Cell Signaling Technology], mouse IgG2a anti-SDHA [sc166947; Santa Cruz Biotechnology]). The following day, cells were washed with PBS several times, and then incubated with one or more of the following secondary antibodies in blocking solution for 30 minutes (Donkey anti-rabbit Alexa 488 [A21206; Fisher Scientific], Donkey anti-mouse Alexa Fluor 488 [A21202; Fisher Scientific], Donkey anti-rabbit Alexa Fluor 555 [A31572; Fisher Scientific], Donkey anti-mouse Alexa Fluor 555 [A31570; Fisher Scientific], Donkey anti-rabbit Alexa Fluor Plus 647 [PIA32795; ThermoFisher Scientific], Goat anti-mouse IgG1 Alexa Fluor 488 [A21121; Fisher Scientific], Goat anti-mouse IgG2a Alexa Fluor 555 [A21137; Fisher Scientific], or Donkey anti-rabbit Alexa Fluor Plus 647 [PIA32795; ThermoFisher Scientific]). Where indicated, cells were then stained with a 1:1000 dilution of DAPI (62248; ThermoFisher Scientific) in PBS for 10 minutes. Cells were then washed with PBS several times prior to imaging.

### Confocal microscopy

Except where indicated below for cell migration analysis, all images were acquired on a Nikon Ti2 microscope equipped with a Yokogawa CSU-W1 spinning disk confocal, a Hamamatsu Orca-Fusion sCMOS camera, and a Nikon 100x (1.45 NA) objective. For live cell imaging, cells were maintained in an environmental chamber and incubated at 37°C, 5% CO_2_. All images were acquired using Nikon Elements. All z-series images were acquired with a 0.2 µm-step size. Image analysis (detailed below), maximum intensity projections, and other image adjustments were made with ImageJ/Fiji. Linescans were generated using a slightly modified version of the RGB Profiles Tool macro (https://imagej.nih.gov/ij/macros/tools/RGBProfilesTool.txt) in ImageJ/Fiji.

### Immunofluorescence analysis and quantification

#### METEOR identification and Fluorescence Intensity Ratio quantification

Fixed cells were co-immunolabeled as described above with a control MICOS antibody (mouse anti-MIC60 or rabbit anti-MIC19 labeled with a secondary antibody conjugated to Alexa Fluor 488 (AF488)) along with a primary antibody targeting a query protein labeled with a secondary antibody conjugated to Alexa Fluor 555 (AF555). Z-series images were acquired for at least two independent experiments for each antibody, and maximum intensity projection images were analyzed. Mitochondria that were qualitatively MICOS-enriched were identified in AF488-labeled images blind to the AF555 signal for the query protein. Regions of interest (ROIs) tightly cropped to the MICOS-AF488 enriched-site (ROI_METEOR_) and a corresponding well-resolved mitochondria in the cell periphery (ROI_periphery_) were identified. The maximum fluorescence intensities of the AF555 signal in the ROI_METEOR_ and ROI_periphery_ were determined, and the ratio between the two was computed to determine the Fluorescence Intensity Ratio of the query protein. The number of METEORs analyzed for each antibody is displayed in the corresponding figure legends. To quantify MIC60-AF555 enrichment, MIC19-AF488 was used to blindly identify METEORs.

The Fluorescence Intensity Ratio of MIC60 at METEORs was 2-fold or greater at 95% of sites (see Figure 1E). Therefore, we used a cutoff of 2-fold MIC60 enrichment to define METEOR sites in subsequent experiments, including determining the number of METEORs per cell, determining the proximity of METEORs to the PM, determining whether mitochondria in filopodia were METEORs, and assessing the presence of mtDNA. To exclude examples of non-specific MIC60 labeling, METEORs were operationally defined as positive for a second mitochondrial protein (i.e. TIMM23). Image analysis of knockdown or treated cells was always performed blind to sample identity.

### Analysis of filopodia length, density, and presence of mitochondria

The percentage of filopodia which contain mitochondria was determined by labeling cells with WGA-CF405 and immunolabeling TIMM23 and MIC60, and manually counting the filopodia in each cell and the number of filopodia that contain mitochondria. The length and density of filopodia was measured manually with ImageJ/Fiji. To determine filopodia density, the number of filopodia relative to the cell perimeter measured in microns was determined for the portion of individual cells that were not in contact with other cells. To compare the length of filopodia containing mitochondria compared to those absent of mitochondria, the length of an immediately adjacent filopodium was measured. In all cases, identity of images was blinded prior to analysis.

### Analysis of mitochondrial distribution and morphology

To determine mitochondrial distribution in control versus MYO19-depleted cells, we examined cells stained with WGA-CF405S and immunolabeled with TIMM23. ImageJ/Fiji was used to manually calculate the distance from the center of the nucleus to the most peripheral mitochondrion in that cell, and the distance from the same point in the nucleus to the position of the PM nearest to the mitochondrion. The ratio between the two distances was then calculated for the indicated number of individual cells per condition as described in legends. To determine mitochondrial morphology, images were categorized as indicated in legends. In the case of both measurements, sample identity was blinded prior to analysis.

### Analysis of mitochondrial membrane potential

For analysis of mitochondrial membrane potential, cells were first transiently transfected with PM-TagBFP as described above. Adherent cells were treated with 200 nM MitoTracker Green (ThermoFisher Scientific M7514) for 30 minutes, washed with complete media, and then treated with 50-100 nM TMRE (ThermoFisher Scientific T669) for 15 minutes. Cells were washed and media was replaced with Fluorobrite DMEM (ThermoFisher Scientific A1896701) supplemented with 10% FBS prior to imaging by confocal microscopy. To determine the fluorescence ratio of TMRE in filopodia versus cell peripheral mitochondria, maximum intensity projections were made, excluding image planes with non-specific staining of the microscope dish. Mitochondria in filopodia and the cell periphery were identified with MitoTracker Green, and the ratio of the background-subtracted maximum fluorescence intensity of the TMRE staining at each mitochondrion was determined. As a control, PM-TagBFP transfected cells were labeled as described above and imaged prior to and after 45 minutes treatment with 10 µM CCCP (Sigma-Aldrich C2759). Individual mitochondria in the cell periphery were identified with MitoTracker Green and the ratio of the background-subtracted maximum fluorescence intensity of TMRE signal post-treatment relative to pre-treatment was determined.

### Electron microscopy and analysis of cristae density

After treatment with control siRNA or MYO19-targeted siRNA as described above, ∼50,000 cells were allowed to adhere to poly-D-lysine coated glass-bottom microscope dishes (MatTek P35GC-1.5-14-C) for 24 hours. Cells were fixed with 2.5% glutaraldehyde in 0.1 M sodium cacodylate buffer and submitted to the UTSW Electron Microscopy Core Facility for processing as described previously (*28*). Briefly, after five rinses in 0.1 M sodium cacodylate buffer, cells were post-fixed in 1% osmium tetroxide and 0.8 % K_3_[Fe(CN)_6_] in 0.1 M sodium cacodylate buffer for 1h at 4°C. Cells were kept overnight at 4°C in water, and then en bloc stained with 2% aqueous uranyl acetate in the dark at room temperature. After five rinses with water, specimens were dehydrated with increasing concentration of ethanol at room temperature, infiltrated with a ratio of 2:1 ethanol to Embed-812 resin, followed by a 1:2 solution of ethanol to resin, followed by pure resin, and then polymerized in a 60°C oven overnight. Embed-812 discs were removed from MatTek plastic housing by submerging the dish in liquid nitrogen. Pieces of the disc were glued to blanks with super glue and blocks were sectioned with a diamond knife (Diatome) on a Leica Ultracut UCT (7) ultramicrotome (Leica Microsystems) and collected onto copper grids and post-stained with 2% uranyl acetate in water and lead citrate. Images were acquired on a JEM-1400 Plus transmission electron microscope equipped with a LaB6 source operated at 120 kV using an AMT-BioSprint 16M CCD camera. Images were blinded post-acquisition prior to analysis. Approximately 3-5 well-resolved mitochondrial profiles were analyzed per tomograph, and 3-5 tomographs were analyzed per cell. The ratio of the number of cristae per mitochondrial profile to the perimeter of the mitochondrial profile was determined using Fiji/ImageJ and manually computed.

### Analysis of cell migration

For all cell migration assays, cells were treated as indicated and ∼6,000 cells were seeded per well of an 8-well µ-slide high glass bottom chamber dish (ibidi 80807) and allowed to adhere overnight. Where indicated, each well was first coated with 300 µL of 0.05 mg/mL collagen I for 1 hour. Cells were then treated with Hoechst dye diluted 1:40,000 in complete media and imaged immediately with a Nikon Eclipse Ti epifluorescence microscope equipped with an ANDOR Zyla sCMOS camera, a Nikon 20x (0.75 NA) air objective, and an environmental chamber (37°C, 5% CO_2_). Single plane images were acquired at an interval of 5 minutes, for 210 minutes, using Micro-Manager. Images were analyzed using the batch and track packages from IMARIS 10.1.0 (Oxford Instruments). Cells were identified from background-subtracted Hoechst images using an estimated diameter of 11 µm. A “quality threshold” was set at approximately 10% to remove non-specific staining. Images were also manually adjusted, where appropriate, to include cells missed by the algorithm. To track cells, autoregressive motion with a maximum distance of 10 µm per frame and a gap size of 2 frames was used. After analysis, data for displacement length (the distance between the starting and ending positions of a cell) and time since track start (the number of frames a cell was successfully tracked for) were extracted. Displacement rate was calculated by creating a ratio of displacement length over the calculated track duration and was plotted for individual cells as indicated in legends.

### Statistical Analysis

Statistical analyses were performed using Graph Pad Prism 10.1.2. Statistical tests were performed as indicated in figure legends.

## Acknowledgements

We thank Natalie Niemi for critical reading of the manuscript and helpful discussions. We thank Mia Hofstad for technical advice. We thank the UT Southwestern Quantitative Light Microscopy Facility (supported in part by P30CA142543 and equipment grant NIH 1S10OD028630-01) for technical support and access to microscopes. We thank Ethan Ozment and Phoebe Doss in the UT Southwestern Electron Microscopy Core Facility (supported by NIH 1S10OD021685-01A1) for sample preparation and assistance with image acquisition. This work was supported by a grant from the NIH to JF (R35GM137894). The authors declare no competing financial interests.

